# Reduced methionine synthase (*Mtr*) expression creates a functional vitamin B12 deficiency that leads to uracil accumulation in mouse mitochondrial DNA

**DOI:** 10.1101/2022.08.29.505750

**Authors:** Katarina E. Heyden, Joanna L. Fiddler, Yuwen Xiu, Olga V. Malysheva, Michal K. Handzlik, Whitney N. Phinney, Linsey Stiles, Sally S. Stabler, Christian M. Metallo, Marie A. Caudill, Martha S. Field

## Abstract

Adequate thymidylate (dTMP or the “T” base in DNA) levels are essential for stability of mitochondrial DNA (mtDNA) and nuclear DNA (nDNA). Folate and vitamin B12 (B12) are essential cofactors in folate-mediated one carbon metabolism (FOCM), a metabolic network which supports synthesis of nucleotides (including dTMP) and methionine. Perturbations in FOCM impair dTMP synthesis, causing misincorporation of uracil (or a “U” base) into DNA. During B12 deficiency, cellular folate accumulates as 5-methyltetrahdryfolate (5-methyl-THF), limiting nucleotide synthesis. The purpose of this study was to determine how B12 deficiency and dietary folate interact to affect mtDNA integrity and mitochondrial function in mouse liver. Mice expressing reduced methionine synthase (*Mtr*) levels were used to create a functional B12 deficiency. Folate accumulation, uracil levels, mtDNA content, and oxidative phosphorylation capacity were measured in male *Mtr*^*+/+*^ and *Mtr*^*+/-*^ mice weaned onto either a folate-sufficient control diet (2 mg/kg folic acid, C) or a folate-deficient diet (FD, lacking folic acid) for 7 weeks. *Mtr* heterozygosity led to increased liver 5-methyl-THF levels. *Mtr*^*+/-*^ mice consuming the C diet also exhibited a 40-fold increase in uracil in liver mtDNA. However, the combination of *Mtr* heterozygosity and exposure to the FD diet partially alleviated the level of uracil accumulation in mtDNA. Furthermore, *Mtr*^*+/-*^ mice exhibited a 25% decrease in liver mtDNA content and a 20% decrease in maximal oxygen consumption rates. Impairments in mitochondrial FOCM are known to lead to increased uracil in mtDNA. This study demonstrates that impaired cytosolic dTMP synthesis also leads to increased uracil in mtDNA.

## Introduction

Vitamin B12 (B12) deficiency disproportionately impacts older adults, vegans/vegetarians, and pregnant women (1). Hematological and neurological symptoms are typical of B12 deficiency, due to the role of B12 in DNA synthesis. B12 and folate (vitamin B9) are essential cofactors required for folate-mediated one-carbon metabolism (FOCM), a metabolic pathway which provides one-carbon groups for *de novo* biosynthesis of nucleotides and methyl donor generation (1). Only two mammalian enzymes require B12 as a cofactor: methyl malonyl CoA mutase (MCM), which supports branched-chain amino acid metabolism and resides in the mitochondria, and methionine synthase (MTR), which is part of FOCM and localizes to the cytosol (1). MTR uses both folate and B12 co-factors to catalyze regeneration of methionine from homocysteine. In this two-step process, 5-methyl-THF donates a methyl group to cobalamin (a form of B12), releasing THF and generating methylcobalamin (2). Methylcobalamin then provides the methyl group to homocysteine for methionine synthesis (2). The conversion of 5,10-methylene-THF to 5-methyl-THF by methylenetetrahydrofolate reductase (MTHFR) is irreversible, and the only enzyme capable of metabolizing 5-methyl-THF to release THF is MTR (3) (Fig. 1). Without adequate B12 levels or as a result of decreased MTR levels, folate becomes increasingly trapped in the 5-methyl-THF form, which decreases the availability of other folate cofactors (i.e., THF and 5,10-methylene-THF) for key processes such as nucleotide synthesis. Therefore, a hallmark of B12 deficiency is the cellular accumulation of 5-methyl-THF, a phenomenon commonly known as the “5-methyl trap” (3).

**Figure 1.**
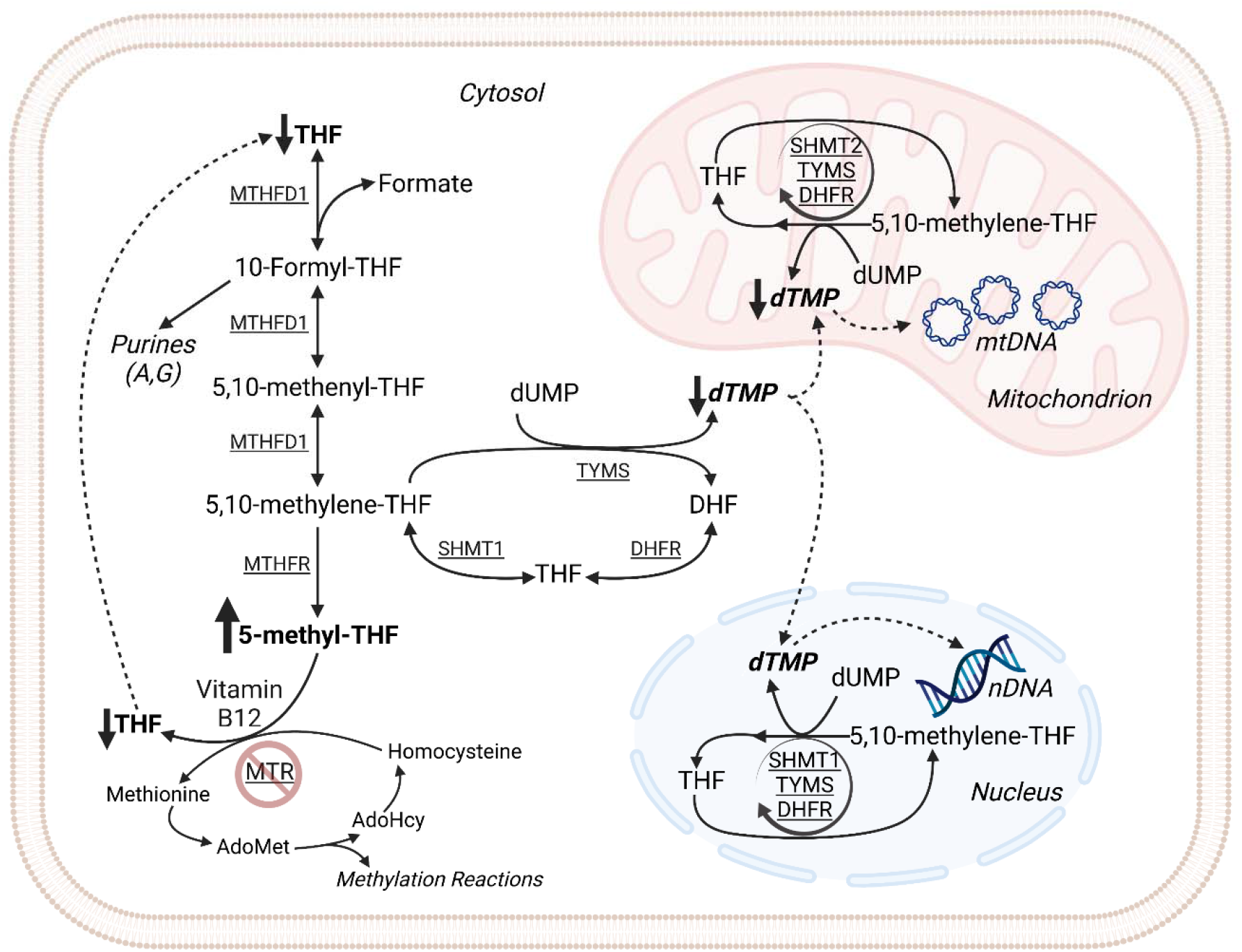
The effects of decreased *Mtr* expression on folate-mediated one-carbon metabolism. Vitamin B12 deficiency, as modeled by partial knockout of the *Mtr* gene in mice causes elevation of cellular folate as 5-methyl-THF and matched reduction of folate as THF. The conversion of 5,10-methylene-THF to 5-methyl-THF is irreversible, and 5-methyl-THF can only be metabolized to THF by the MTR enzyme using B12 as a cofactor. AdoHcy, S-adenosylhomocysteine; AdoMet, S-adenosylmethionine; DHFR, dihydrofolate reductase; dTMP, deoxythymidine monophosphate; dUMP, deoxyuridine monophosphate; FOCM, folate-mediated one-carbon metabolism; MTHFD1, methylenetetrahydrofolate dehydrogenase 1; MTHFR, methylenetetrahydrofolate reductase; MTR, methionine synthase; mtDNA, mitochondrial DNA; nDNA, nuclear DNA; SHMT, serine hydroxymethyltransferase; THF, tetrahydrofolate; TYMS, thymidylate synthase.

As mentioned above, folate is required for synthesis of thymidylate (dTMP, or the “T” base in DNA). dTMP is a precursor of deoxythymidine triphosphate (dTTP). dTTP is unique among DNA bases, as DNA polymerases incorrectly incorporate a uracil (or “U”) base into DNA when dTTP levels are low (4). Uracil misincorporation leads to activation of base-excision repair mechanisms, which in the continued absence of dTMP, lead to DNA double-strand breaks, stalled replication fork progression, and DNA instability (4). Folate-dependent dTMP synthesis occurs in multiple cellular compartments (cytosol, nucleus, and mitochondria) (5, 6). There is evidence that either dietary or genetic FOCM impairments decrease cytosolic and nuclear dTMP synthesis leading to nuclear DNA (nDNA) instability (5).

Mitochondrial DNA (mtDNA) is relatively small (16.5 kb) and circular in structure, making it distinct from nDNA. mtDNA encodes for components of the protein complexes within the mitochondrial membrane that allow for energy production through oxidative phosphorylation (OXPHOS) (7). mtDNA replication is one component of mitochondrial biogenesis, and maintenance of mtDNA is closely tied to mitochondrial function (7). Nucleotide synthesis, specifically *de novo* dTMP synthesis, has been shown to occur in the mitochondria and to be essential for maintaining mtDNA integrity (6). Evidence in cultured cells (8) and mouse liver (9) suggests mtDNA is more sensitive than nDNA to dietary folate deficiency. The effects of impaired B12 deficiency on mtDNA stability are almost completely uncharacterized.

The folate-B12 interrelationship has received increased attention, as observational data in humans has associated the combination of elevated folate status with low or deficient B12 status (known as “high folate/low B12”) with a wide array of negative health outcomes: worsened clinical markers of B12 deficiency, diminished response to B12 treatment, and increased risk for cognitive impairment and decline among the elderly (10, 11). However, there is little data from B12-deficient model systems investigating the molecular mechanisms underlying these associations. This study used genetic disruption of *Mtr* in mice (12) to create a functional B12 deficiency. Decreased *Mtr* expression perturbed cellular folate distribution, increased uracil accumulation in liver mtDNA, decreased liver mtDNA content, and reduced mitochondrial respiratory capacity in liver. Interestingly, the increased uracil in mtDNA was attenuated in *Mtr*^*+/-*^ mice exposed to folate-deficient diets. These data provide one of the first mechanistic explanations for how “high folate/low B12” status may cause adverse physiological outcomes.

## Results

### Reduced *Mtr* expression alters whole liver folate distribution but does not affect total folate concentrations

*Mtr*^*+/+*^ and *Mtr*^*+/-*^ mice were weaned onto either control (C; AIN93G-based diet with 2 mg/kg folic acid) or folate-deficient (FD; AIN93G-based diet lacking folic acid) diet and maintained on these defined diets for seven weeks until sacrifice. Mice on the FD diet had slightly higher overall food intake than those on C diet (*p*<0.01), though there were no differences in mouse body weight between groups (*SI Appendix*, Fig. S1A-B). Liver MTR protein levels were reduced by 60% (*p*< 0.001) in *Mtr*^*+/-*^ mice consuming the C diet (Fig. 2A). Interestingly, liver MTR protein levels were also reduced (*p*< 0.001 for effect of diet) as a result of FD diet exposure (Fig. 2A). The same diet and genotype effects were also observed in kidney tissue (*p*=0.04 and *p*=0.018, respectively; *SI Appendix*, Fig. S2).

**Figure 2.**
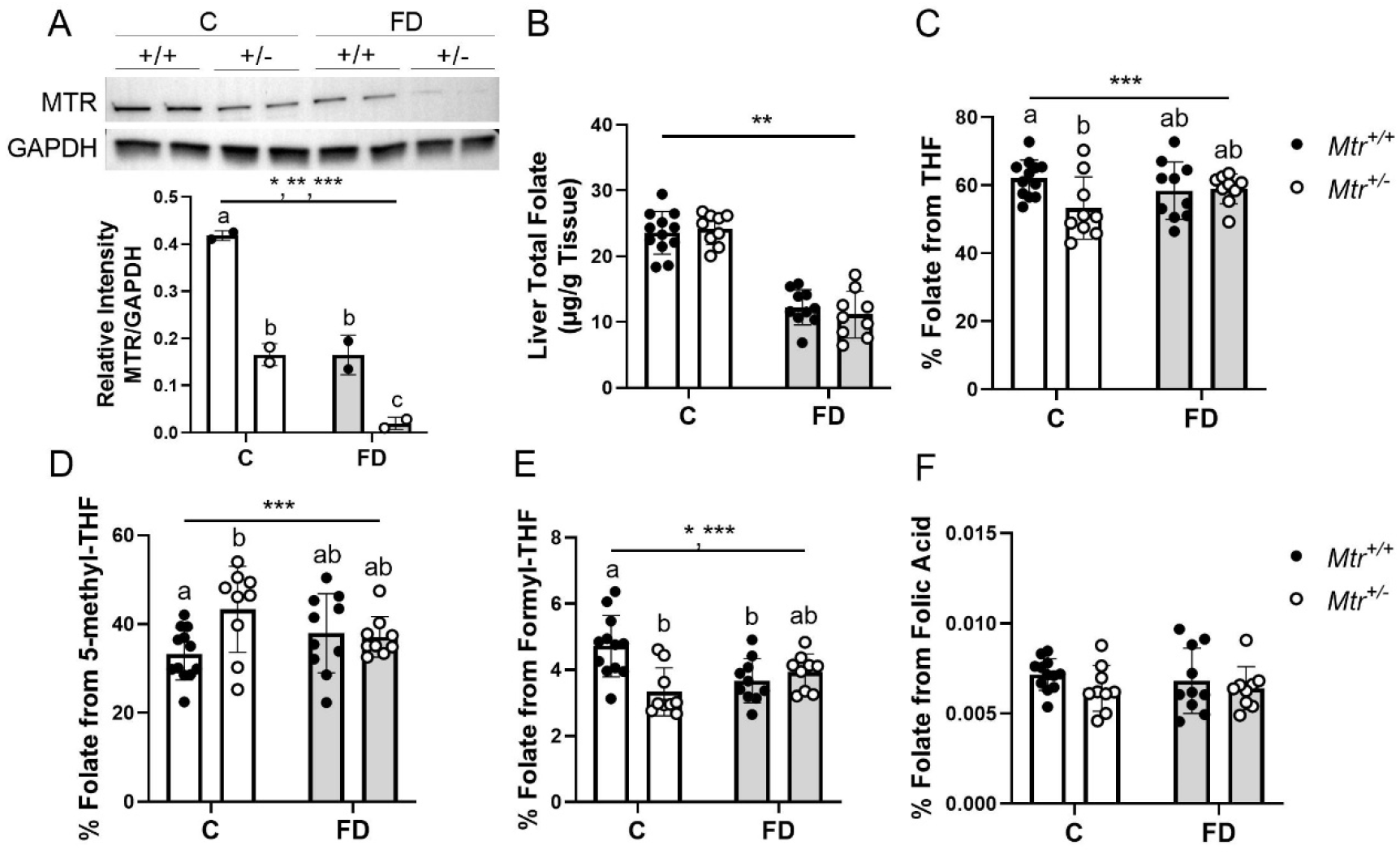
Total folate and folate distribution in *Mtr*^*+/+*^ and *Mtr*^*+/-*^ liver. (A) Liver MTR protein levels normalized to GAPDH; n = 2 per group. (B) Total liver folate; n = 9-12 per group. (C) Liver folate distribution from THF; n = 9-12 per group. (D) Liver folate distribution from 5-methyl-THF; n = 9-12 per group. (E) Liver folate distribution from Formyl-THF; n = 9-12 per group. (F) Liver folate distribution from folic acid; n = 9-12 per group. Folate distribution was quantified using LC-MS/MS. Two-way ANOVA with Tukey’s post-hoc analysis was used to assess main effects of diet and genotype and diet-genotype interaction. Data are presented as mean ± s.d. with statistical significance defined *p* **≤** 0.05. Groups not connected by a common letter are significantly different. * significant genotype effect, ** significant diet effect, *** significant genotype-diet interaction. C, control diet; FD, folate-deficient diet; GAPDH, glyceraldehyde-3-phosphate dehydrogenase; MTR, methionine synthase; THF, tetrahydrofolate.

Folate deficiency and reduced *Mtr* expression elicited changes in plasma transmethylation metabolite levels (Table 1). As previously observed (13), changes in transmethylation biomarkers were observed with FD diet exposure, which increased homocysteine (*p*=0.018), cystathionine (*p*<0.0001), and methylglycine (*p*=0.023), and decreased cysteine (*p*=0.052) and α-aminobutyric acid (*p*=0.02; Table 1). *Mtr*^*+/-*^ genotype-driven changes were observed in methionine and methylglycine levels (*p*=0.046 and *p*<0.001, respectively; Table 1). As expected, plasma folate markedly decreased with FD diet exposure (*p*<0.0001; Table 1). In addition, *Mtr*^*+/-*^ genotype did not impact plasma folate accumulation, in agreement with previous findings in this model (14). As anticipated, plasma MMA was unaffected by either *Mtr*^*+/-*^ genotype or folate deficiency (Table 1), indicating that the *Mtr*^*+/-*^ model of functional B12 deficiency does not affect methylmalonyl-CoA mutase enzyme activity (i.e., the model is specific to the B12-dependent enzymatic activity involved in FOCM).

**Table 1.**
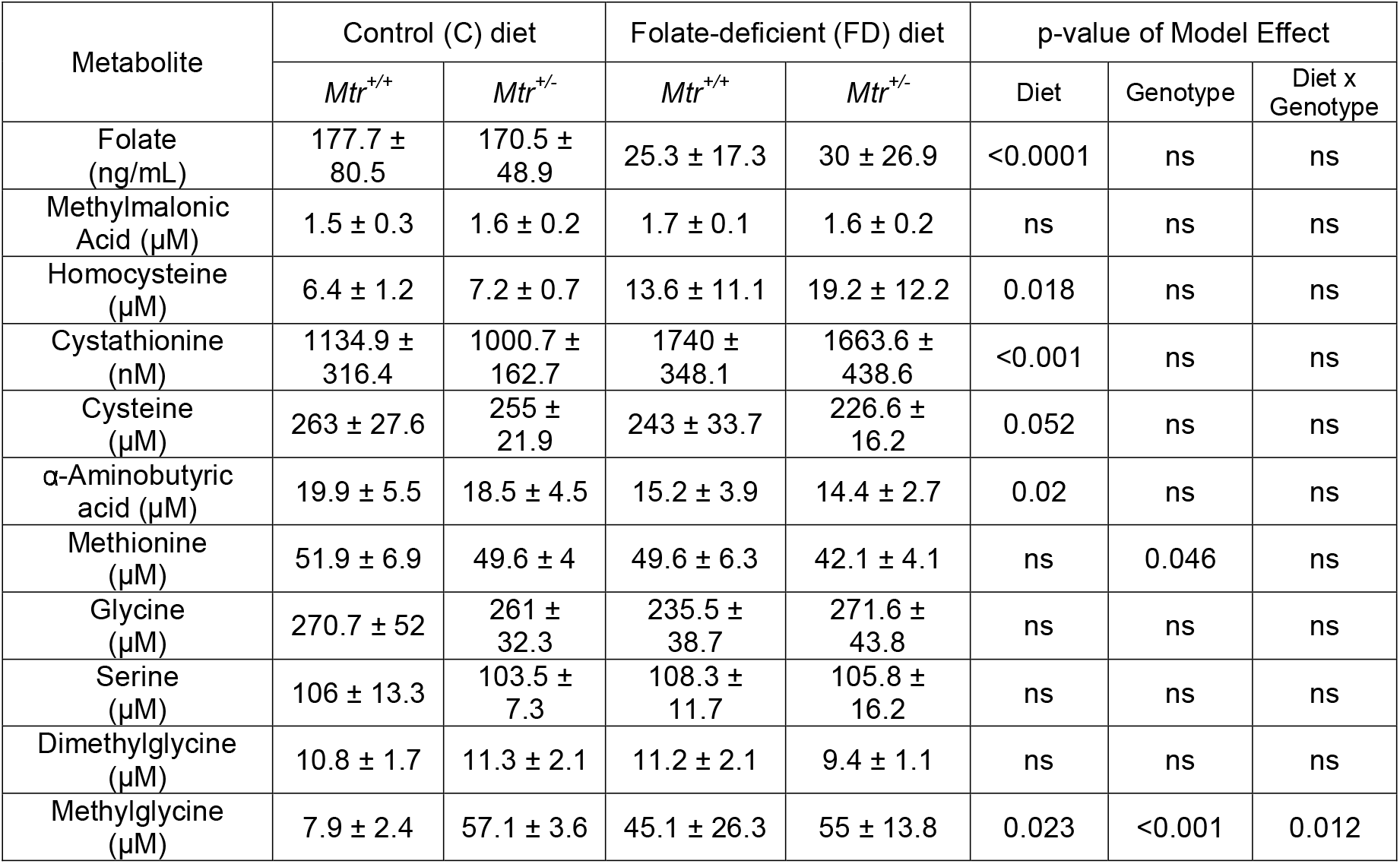
Plasma Metabolite Profile for *Mtr*^*+/+*^ and *Mtr*^*+/-*^ mice. Folate concentration was measured using L. *casei* microbiological assay. MMA concentration was measured by GC-MS. All other biomarkers were measured via stable isotope dilution capillary GC-MS. Data were analyzed by two-way ANOVA with Tukey’s post-hoc analysis for diet and genotype main effects and diet-genotype interaction. Data are presented as mean ± s.d. with statistical significance defined as *p* ≤ 0.05; n = 4-8 per group. ns, not significant.

Exposure to the FD diet reduced total liver folate levels of both *Mtr*^*+/+*^ and *Mtr*^*+/-*^ mice by 50% when measured by LC-MS (*p*<0.0001; Fig. 1B) and microbiological assay (*p*=0.014; *SI Appendix*, Fig. S3). *Mtr* genotype did not influence total liver folate accumulation (Fig. 2B, *SI Appendix*, Fig. S3). One-carbon metabolism consists of several one-carbon substituted folate forms to serve as cofactors and dietary vitamin B12 deficiency is associated with accumulation of folate as 5-methyl-THF; therefore, folate cofactor distribution was quantified. In mice consuming the C diet, reduced *Mtr* expression increased the amount of whole-cell liver 5-methyl-THF from 33% to 43% of total folate (*p*=0.02), with corresponding decreases in the percentage of folates from THF (*p*=0.038) and formyl-THF (*p*<0.001) (Fig. 2C-D). These changes in folate distribution support the presence of a 5-methyl-THF trap in *Mtr*^*+/-*^ whole-cell liver exposed to the C diet. Notably, 5-methyl-THF accumulation was normalized in *Mtr*^*+/-*^ mice consuming the FD diet (Fig. 2C-D) Interestingly, the amount of liver folate in the form of folic acid was very low, accounting for less than 0.01% of all folates even in mice consuming the C diet (Fig. 2F). This suggests that folic acid does not appreciably accumulate in liver of mice consuming defined diets containing 2 mg/kg folic acid.

Total folate levels in liver mitochondria decreased by 20% (*p*=0.03) in mice consuming the FD diet (Fig 3A), as assessed by LC-MS analysis, and confirmed by *L. casei* microbiological assay (*p*=0.01; *SI Appendix*, Fig. S3). As with whole-cell liver folate levels, *Mtr* genotype did not impact total folate levels in liver mitochondria (Fig. 3A). The primary form of folate detected in liver mitochondrial fractions was THF, which consisted of >90% of mitochondrial folates (Fig. 3B) and was consistent with previous literature (15, 16). While *Mtr*^*+/-*^ genotype increased 5-methylTHF levels in liver mitochondria of mice on C diet (*p*<0.0001, Fig. 3C), this increase from 2% to 4% may not be biologically meaningful. There was no detectable level of folic acid present in liver mitochondria.

**Figure 3.**
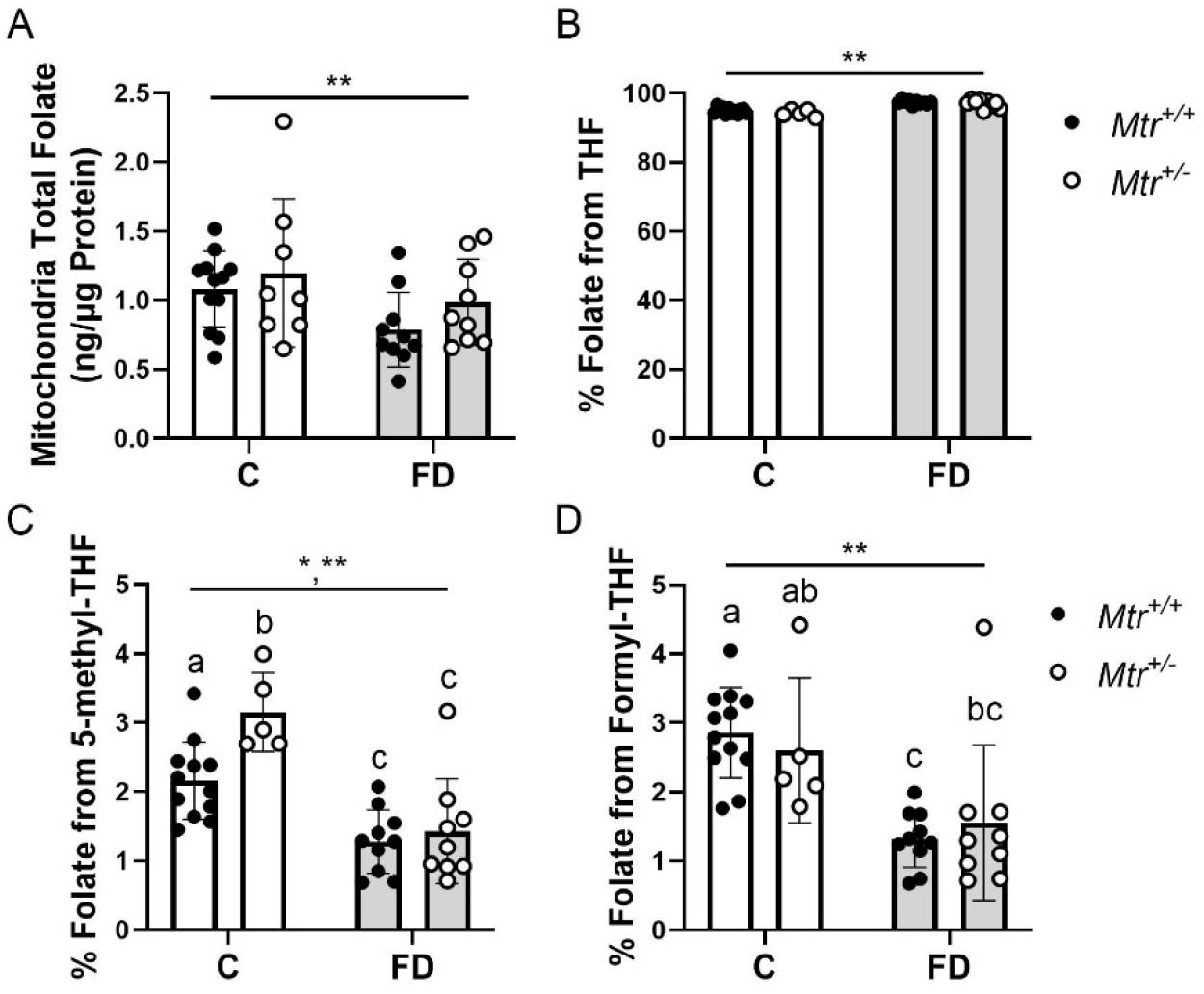
Total folate and folate distribution in mitochondrial fraction from *Mtr*^*+/+*^ and *Mtr*^*+/-*^ liver. (A) Total mitochondrial folate; n = 9-12 per group. (B) Folate distribution from THF; n = 9-12 per group. (C) Folate distribution from 5-methyl-THF; n = 9-12 per group. (D) Folate distribution from 5-formyl-THF; n = 9-12 per group. Folate distribution was quantified using LC-MS/MS. Data are shown as mean ± s.d.. Two-way ANOVA with Tukey’s post-hoc analysis was used to assess main effects of diet and genotype and diet-genotype interaction with statistical significance defined as *p* ≤ 0.05. Groups not connected by a common letter are significantly different.19ignificantant genotype effect, ** significant diet effect, *** significant genotype-diet interaction effect. C, control diet; FD, folate-deficient diet; Mtr, methionine synthase; THF, tetrahydrofolate.

### *Mtr* genotype leads to uracil misincorporation in mtDNA and impaired mitochondrial function

We recently developed an assay to quantify uracil misincorporation into mtDNA (9). As described in detail above, uracil misincorporation arises when *de novo* dTMP synthesis is impaired by either folate or B12 deficiency (5, 17). To assess whether the altered whole-cell folate distribution with *Mtr* loss could affect mtDNA integrity, uracil content of liver mtDNA was measured by quantitative real-time PCR assay (9). In liver mitochondria, there was a significant interaction between diet and genotype for mtDNA uracil accumulation (*p*=0.0001, Fig 4A). Compared to *Mtr*^*+/+*^ mice consuming the C diet, both reduced *Mtr* expression (*p*<0.001) and folate deficiency (*p*<0.001) were associated with increased uracil accumulation in mtDNA (Fig. 4A). In fact, the *Mtr*^*+/-*^ genotype increased uracil levels in mtDNA by nearly 40-fold compared to *Mtr*^*+/+*^ genotype in mice consuming the C diet (Fig. 4A). However, when *Mtr* loss was combined with FD diet, uracil accumulation was attenuated (*p*=0.0001, Fig. 4A), suggesting a protective effect against uracil accumulation in mtDNA in this experimental group. As observed previously (9), uracil was enriched in specific regions of mtDNA (*SI Appendix*, Fig. S4). Importantly, uracil accumulation observed in liver mtDNA with reduced *Mtr* expression or exposure to the FD diet was not observed in liver nDNA (*SI Appendix*, Fig. S5), as observed in other models (9). These data suggest that the 5-methyl-THF trap in whole cell liver causes a disturbance in the cytosolic folate pool and leads to increased uracil misincorporation to mtDNA in mice consuming adequate dietary folate (C diet). Exposure to the FD diet attenuates both the whole-cell 5-methyl-THF accumulation (Fig. 2D) and uracil accumulation in mtDNA (Fig. 4A), suggesting that the decrease in whole-cell 5-methyl-THF levels induced by FD diet also decreases uracil in mtDNA.

**Figure 4.**
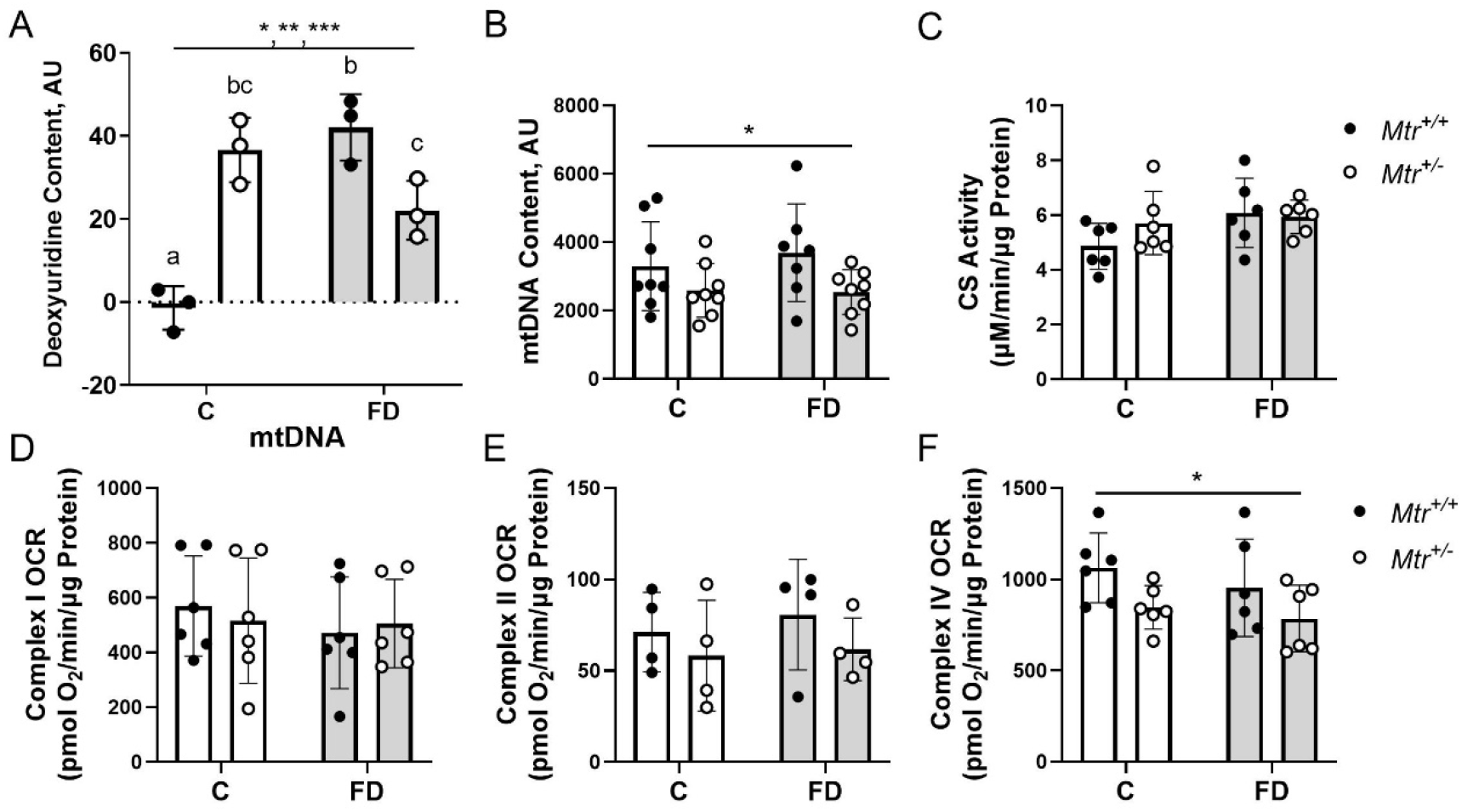
Biomarkers of impaired mitochondrial one-carbon metabolism. (A) Uracil content in liver mitochondrial DNA; n = 3 per group. (B) mtDNA copy number in liver determined using real-time PCR; n = 6 per group. (C) Mitochondrial mass quantified using citrate synthase activity in liver; n = 6 per group. Data are shown as mean ± s.d. and were analyzed by two-way ANOVA with Tukey’s post-hoc analysis, with significance defined as *p* ≤ 0.05. Groups not connected by a common letter are significantly different. * significant genotype effect, ** significant diet effect, *** significant genotype-diet interaction effect. AU, arbitrary units; C, control diet; CS, citrate synthase; FD, folate-deficient diet; Mtr, methionine synthase; OCR, oxygen consumption rate.

In addition to investigating the integrity of mtDNA, we also quantified mtDNA content, which is increasingly recognized as an indicator of mitochondrial dysfunction (7, 18). Reduced *Mtr* expression was associated with a 25% decrease in mtDNA content in mouse liver (*p*=0.045, Fig. 4B). This observed decrease was not due to a decrease in mitochondrial mass in *Mtr*^*+/-*^ liver samples, as there was no difference in citrate synthase activity between groups (Fig. 4C).

To understand the effects of reduced *Mtr* expression on mitochondrial oxidative phosphorylation, maximal respiratory complex activity was measured in liver. There were no changes in activity of complex I or complex II with *Mtr* loss or FD diet exposure (Fig. 4D-E). Interestingly, decreased *Mtr* expression decreased the activity of complex IV by 20% (*p*=0.026, Fig. 4F). These findings suggest that a functional B12 deficiency specific to FOCM impairs mitochondrial oxidative phosphorylation capacity.

## Discussion

In this study, the *Mtr*^*+/-*^ mouse model was used to assess whether a functional vitamin B12 deficiency (induced by reduced *Mtr* expression) affected mtDNA integrity and mitochondrial function. *Mtr*^*+/-*^ mice exhibit decreased MTR protein levels in liver (Fig. 1A) and have been previously shown to be a model for functional B12 deficiency (12). As mentioned earlier, B12 is required by only two enzymes: MTR and MCM. Elevated MMA, a byproduct of excess MCM substrate, is a classical marker to diagnose B12 deficiency. Notably, the *Mtr*^*+/-*^ mouse model does not affect MMA levels (Table 1), meaning that the model isolates the effects of B12 deficiency to those effects coming from impaired FOCM (Fig 6).

As expected, decreased *Mtr* expression shifted liver whole-cell folate distribution to cause an increase in 5-methyl-THF levels at the expense of THF levels (Fig 2C-D). The majority of total cellular folate is distributed evenly between the cytosolic and mitochondrial compartments (with about 10% in the nuclear compartment) (19), and given that reduced *Mtr* expression did not notably alter mitochondrial folate distribution (Fig. 3B-D), the changes observed in whole-cell folate distribution are likely reporting on the accumulation of 5-methyl-THF in the cytosol. Thus, reduced *Mtr* expression causes a cytosolic “5-methyl-trap” in mouse liver, or functional B12 deficiency, even in the presence of sufficient dietary vitamin B12 levels.

As described earlier, an accumulation of 5-methyl-THF decreases folate cofactors needed for nucleotide biosynthesis, impairing *de novo* dTMP synthesis and DNA replication and/or repair (3). The observation that uracil in mtDNA is elevated in *Mtr*^*+/-*^ mice (Fig. 4A) suggests that the mitochondrial compartment is sensitive to changes in cytosolic *de novo* dTMP synthesis capacity and/or that a cytosolic 5-methyl-trap decreases the amount of cytosolic dTMP available for transport to the mitochondria. Mitochondrial dTMP synthesis capacity has been shown to be insufficient to maintain adequate mitochondrial dTMP levels and prevent mtDNA uracil misincorporation in cell culture models (8). In addition, 5-methyl-THF is a tight-binding inhibitor of the enzyme serine hydroxymethyltransferase 1 (SHMT1), which is essential for *de novo* dTMP synthesis (Fig. 1) (5, 20), providing a second mechanism whereby 5-methylTHF accumulation may impair cytosolic dTMP synthesis contributing to uracil accumulation in mtDNA (Fig. 3A).

The phenotype of uracil accumulation in this model was specific to mtDNA, as uracil levels in nDNA remained unchanged (*SI Appendix*, Fig S5.), consistent with our previous findings in the *Shmt2*^*+/-*^ mouse model (9). Taken together, these data suggest that mtDNA is more sensitive to impaired FOCM than is nDNA or that mtDNA uracil may be an earlier biomarker of FOCM dysfunction. In addition, uracil levels in nDNA vary widely by tissue type (5). Studies are needed to determine whether mtDNA uracil levels also vary by tissue type and whether uracil can be detected in lymphocyte DNA, which would facilitate its use as a biomarker.

The physiological consequences of uracil accumulation in mtDNA have not yet been fully investigated. Decreased *Shmt2* expression impairs mitochondrial FOCM and also causes uracil accumulation in liver mtDNA (9). Decreased *Shmt2* did not affect liver mtDNA content or liver mitochondrial mass (9). Importantly, uracil accumulation in the *Shmt2*^*+/-*^ model closely paralleled impairments in oxidative phosphorylation and mitochondrial membrane potential (9). Uracil accumulation in mtDNA as a result of disrupted cytosolic FOCM (i.e. decreased *Mtr* expression) (Fig. 4A) was also associated with impaired maximal oxidative phosphorylation capacity in liver (Fig. 4F). However, reduced *Mtr* expression also led to decreased mtDNA content in liver (Fig. 4B), suggesting an additional means by which *Mtr*^*+/-*^ genotype may perturb mitochondrial function.

As described above, the combination of elevated folate status with low or deficient B12 status is associated with a wide array of negative health outcomes, though mechanisms have not been thoroughly investigated in model systems (10). Notably, this study captures the first evidence of an adverse molecular phenotype (i.e,. increased uracil accumulation in mtDNA) resulting from the combination of “high folate/low B12” status, as mtDNA uracil levels were attenuated in *Mtr*^*+/-*^ mice consuming the FD diet. This finding is compelling because mitochondrial function changes are associated with many of the pathologies identified in the observational literature to be exacerbated as a result of “high folate/low B12” status. Further studies using a more extensive range of dietary folic acid content are needed to more fully elucidate these mechanisms and to determine the utility of mtDNA uracil as a biomarker of risk for adverse pathologies.

## Materials and Methods

### Breeding of *Mtr*^*+/+*^ and *Mtr*^*+/-*^ Mice

The *Mtr*^*+/-*^ mouse model, generated as previously described (12), was backcrossed for more than 10 generations to C57Bl/6J. To characterize the interaction between reduced *Mtr* expression and varying levels of dietary folic acid exposure, C57BL/6J females were intercrossed with *Mtr*^+/-^ males. Male *Mtr*^+/+^ and *Mtr*^+/-^ offspring were then weaned and randomly assigned to one of two defined diets at 3 weeks of age. The diets consisted of defined AIN93G control (C) diet containing 2 mg/kg folic acid (#117814GI; Dyets, Inc., Bethlehem, PA), or an AIN93G-based folate-deficient (FD) diet lacking folic acid (#117815GI; Dyets, Inc., Bethlehem, PA). Dietary intake and body weights were recorded every 14 days. Mice were maintained on these diets for 7 weeks and sacrificed via cervical dislocation following CO_2_ euthanasia. Mice were fasted for 12 hours prior to harvest. Whole blood was collected via cardiac puncture into Heparin-coated tubes, and plasma and red blood cells were separated by centrifugation at 2500 x g in a microcentrifuge tube and immediately flash frozen in liquid nitrogen. Tissues were harvested, rinsed in ice-cold 1X phosphate buffered saline (PBS, Corning), and immediately flash frozen in liquid nitrogen or used for liver mitochondrial isolation (described below). Plasma, red blood cells, and tissues were then stored at −80°C for further analysis.

### Mitochondrial Isolation

Mitochondria were isolated from fresh mouse liver as previously described (9). Mitochondria were saved as pellets or resuspended in extraction buffer (2% (wt/vol) sodium ascorbate, 0.2 M beta-mercaptoethanol, 0.05 M HEPES pH 7.85, 0.05 M CHES pH 7.85) to prevent folate degradation. Samples were stored at −80°C.

### Folate Distribution

Folate levels and vitamer distribution for liver and mitochondrial samples were quantified by LC-electrospray tandem MS adapted from previously described methods (21–24). Liver and mitochondrial samples were weighed at the time of harvest. Liver total folate levels were normalized to liver weight. Mitochondrial total folate levels were normalized to protein concentration, which was assessed by Lowry-Bensadoun Assay (25).

### *Lactobacillus casei* Assay for Total Folates

Liver tissue, liver mitochondria (stored in extraction buffer), and plasma folate concentrations were measured by the microbiological *L. casei* assay as previously described (26). L. casei growth was quantified at 550 nm by Epoch Microplate Spectrophotometer (Biotek Instruments). Total folate measurements for liver and purified liver mitochondria were normalized to protein concentrations for each sample.

### Uracil in mtDNA measured by real-time PCR

DNA was isolated from frozen mitochondrial pellets using a QIAprep Spin Miniprep Kit (Qiagen). Uracil in mitochondrial DNA was quantified using a previously described real-time PCR assay (9).

### Uracil in nuclear DNA measured using gas chromatography-mass spectrometry

DNA was isolated using High Pure PCR Template Preparation Kit (Roche). Following RNAse A treatment, DNA was purified using High Pure PCR Purification Kit (Roche) and concentrations were quantified by Qubit (Thermo Scientific). 2 ug of DNA was treated with uracil DNA glycosylase (UDG, New England Biolabs, Inc.) for 60 minutes at 37°C with gentle shaking. Samples were derivatized and uracil levels quantified as previously described (9).

### Mitochondrial DNA Copy Number qPCR

DNA from liver and cell pellets was isolated using High Pure PCR Template Preparation Kit (Roche). DNA concentrations were measured by Nanodrop 2000c Spectrophotometer (Thermo Scientific). Quantitative PCR to measure mitochondrial DNA content was performed as previously described (27).

### Mitochondrial Mass

Mitochondrial mass in whole liver was measured according to the manufacturer’s instructions using the Citrate Synthase Activity Assay Kit (Sigma) and normalized to protein concentration. Protein concentrations were assessed by Lowry-Bensadoun assay (25). Absorbance of the colorimetric assay was quantified by Epoch Microplate Spectrophotometer (Biotek Instruments).

### Respirometry in frozen samples

The oxygen consumption rate (OCR) of liver mitochondria was measured using the Seahorse XFe24 Extracellular Flux Analyzer (Agilent Technologies) as previously described (28) with minor modifications. Briefly, mitochondria from frozen liver were isolated in ice-cold MAS buffer using a Potter-Elvehjem (Teflon-glass) homogenizer with 10 strokes followed by centrifugation. Protein concentrations of the liver mitochondria supernatant were determined by a Pierce BCA Protein Assay (ThermoFisher). Mitochondrial homogenates (150 µg) were loaded into the assay plates and centrifuged at 2000 g for 5 min at 4°C with no brake. The OCR of each complex was determined using Wave software (Agilent) and was defined by the highest respiratory capacity value following the injection of the corresponding complex stimulating substrate: Complex I, NADH; Complex II, succinate, and Complex IV, TMPD.

### Plasma Metabolite Analysis

Total plasma folate was assessed by *L. casei* assay as mentioned above. Total plasma levels of homocysteine, cystathionine, cysteine, α-aminobutyric acid, methionine, glycine, serine, dimethylglycine, and methylglycine were quantified by stable isotope dilution capillary GC-MS as previously described (29, 30). MMA quantification was performed using 3 uL of plasma that was spiked in with [U-13C]-MMA (Cambridge Isotope Laboratories, cat# CLM-9426-PK) as previously described (31) using gas chromatography mass spectrometry

### Immunoblotting

Tissues were lysed by sonication in lysis buffer (150 mM NaCl, 10 mM Tris-Cl, 5 mM EDTA pH 8, 5 mM dithiothreitol, 1% Triton X-100, protease inhibitor) and cell debris was removed by centrifugation at 4°C for 10 minutes at 14,000 x g. Protein concentrations of tissue or cell lysate supernatants were assessed by Lowry-Bensadoun assay (25). Samples were boiled with SDS-PAGE sample loading buffer (6X SDS) and 30 ug of protein was loaded to each well of a 10% Tris-glycine SDS-PAGE gel (BioRad). Proteins were transferred to a PVDF membrane (Millipore). Membranes were blocked with 5% (wt/vol) non-fat dry milk in 1XPBS with 0.05% Tween-20 for 1 hour. Membranes were then incubated for 1 hour with 1:2000 primary antibody (α-SHMT2 and GAPDH, Cell Signaling #33443S and #14C10; α-MTR, ProteinTech #25896) in 5% bovine serum albumin (BSA) 0.02% NaN_3_Membranes were then washed 3 times in 1XPBS with 0.01% Tween-20 and incubated for 1 hour with HRP-conjugated secondary antibody diluted 1:20,000 in 5% (wt/vol) non-fat dry milk in 1XPBS with 0.05% Tween-20. Membranes were exposed with chemiluminescent substrate (BioRad) and visualized using a ProteinSimple Imager (Bio-techne). Protein levels were quantified using ImageJ software.

### Statistical Analyses

All statistical analyses were performed with R statistical software (version 4.0.4 “Lost Library Book”). Experiments comparing two genotypes (*Mtr*^*+/+, +/-*^) and two different diets (C/FD) were analyzed using a two-way ANOVA with fixed effects of genotype, diet, and genotype-diet interaction, and were followed by Tukey’s post-hoc analysis. Body weight and food intake datasets were analyzed using a two-way ANOVA with mixed effects of diet and time. Model assumptions of normality and homogenous variance for each dataset were confirmed by QQ plot and Levene’s Test, respectively. Data are presented as means ± standard deviation.

## Acknowledgments

This work was supported in part by a President’s Council of Cornell Women grant.

