## Supplementary material for "Reduced methionine synthase (*Mtr*) expression creates a functional vitamin B12 deficiency that leads to uracil accumulation in mouse mitochondrial DNA"

^1^Division of Nutritional Sciences, Cornell University, Ithaca, NY, USA; ^2^Department of Bioengineering, University of California, La Jolla, San Diego, CA, USA; ^3^Department of Medicine, University of Colorado, Aurora, CO, USA; ^4^Department of Medicine/Division of Endocrinology, University of California, Los Angeles, CA USA; ^5^Molecular and Cell Biology Laboratory, Salk Institute for Biological Studies, La Jolla, CA, USA

Corresponding Author: Martha S. Field

**This file includes:**

Figures S1 to S5

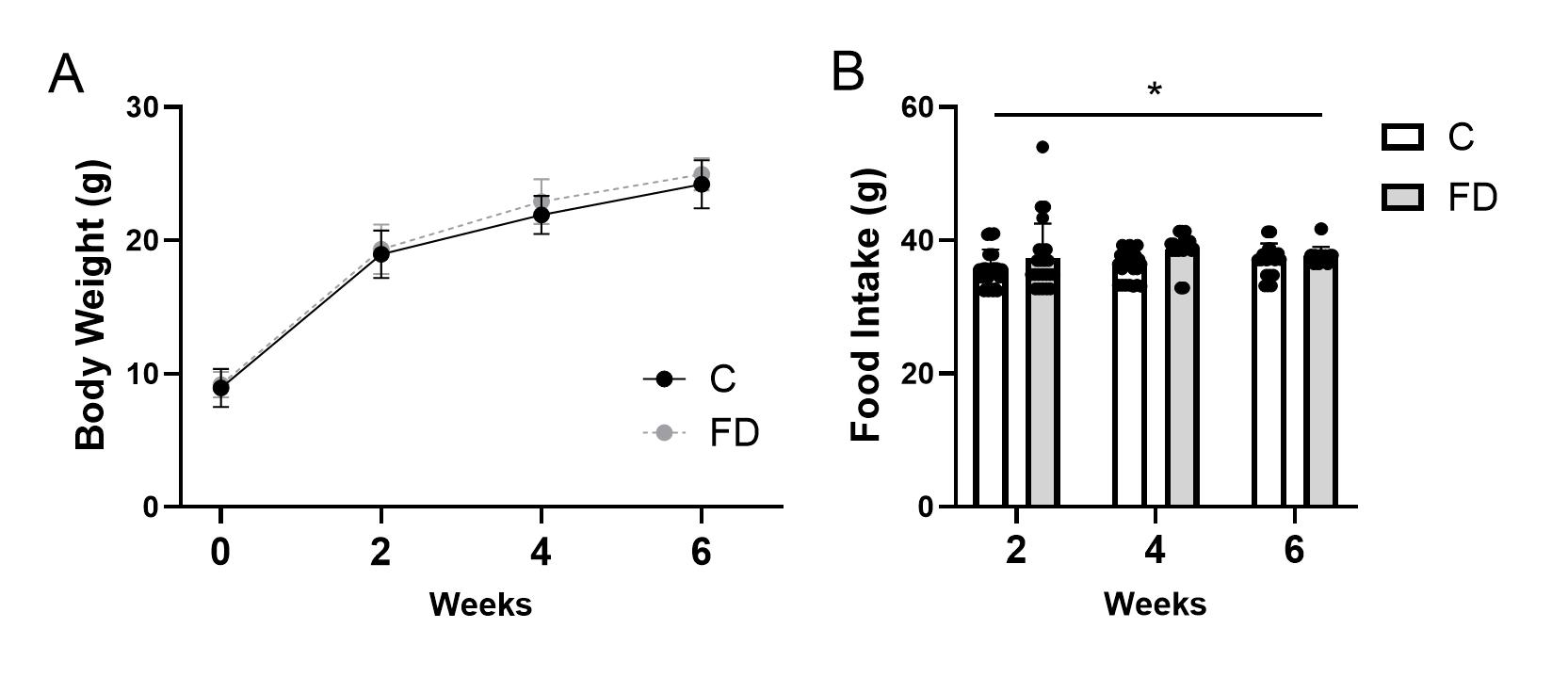

Fig. S1. Body weight and food intake measurements. At weaning, mice were placed on the C or FD diet and (A) body weights and (B) food intake were measured at weaning and every two weeks thereafter; n = 20-25 per group. Two-way mixed ANOVA was used to assess fixed effects of genotype, diet, and genotype-diet interaction. Residual analyses were performed to check the model assumptions of normality and homogeneous variance. Data are shown as mean ± s.d. and significance was defined as p ≤ 0.05. *significant genotype effect, ** significant diet effect, *** significant genotype-diet interaction effect. C, control diet; FD, folate-deficient diet.

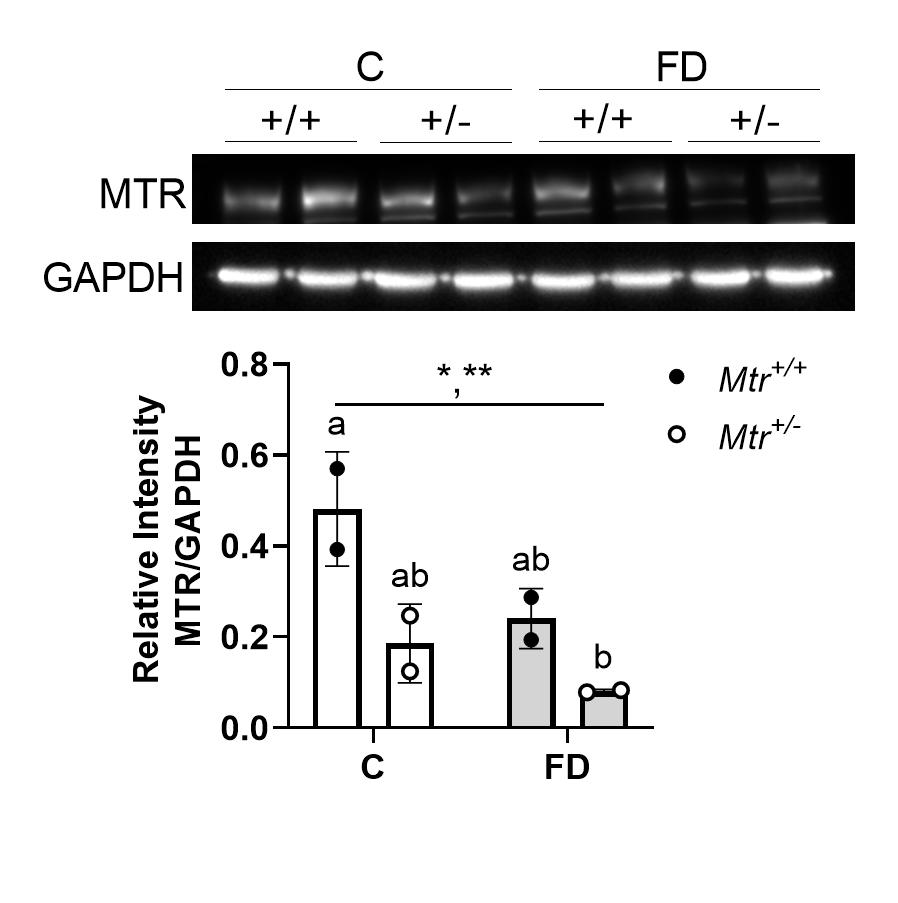

Fig. S2. Methionine synthase protein expression in mouse kidney decreases with *Mtr* loss and with folate deficiency. Kidney MTR protein levels normalized to GAPDH; n = 2 per group. Data are shown as mean ± s.d. and were analyzed by two-way ANOVA with Tukey’s post-hoc analysis, with significance defined as *p* ≤ 0.05. Groups not connected by a common letter are significantly different. * significant genotype effect, ** significant diet effect, *** significant genotype-diet interaction effect. C, control diet; FD, folate deficient diet; MTR, methionine synthase; GAPDH, glyceraldehyde-3-phosphate dehydrogenase.

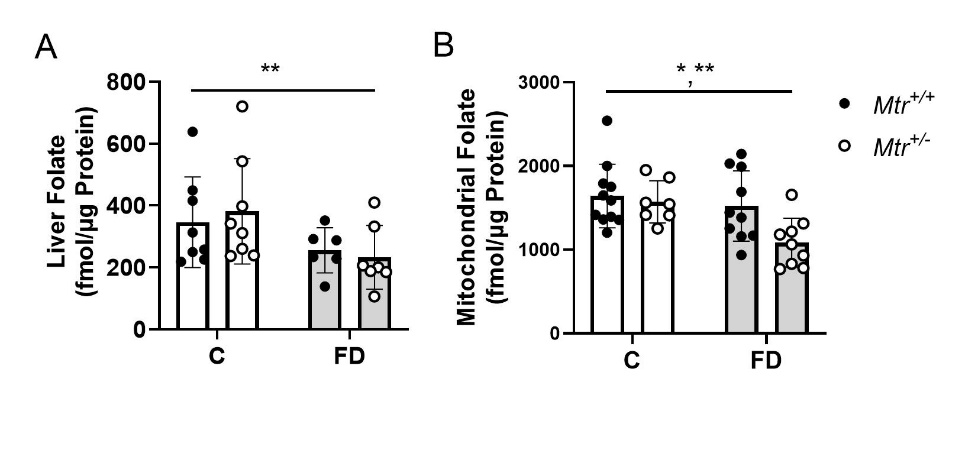

Figure S3. Total folate levels in whole cell and mitochondrial fraction of *Mtr^+/+^* and *Mtr^+/-^* mouse liver. (A) Total whole-cell and (B) mitochondrial folate levels in liver measured by L. *casei* microbiological assay; n = 7-11 per group. Data are shown as mean ± s.d. and were analyzed by two-way ANOVA with Tukey’s post-hoc analysis, with significance defined as *p* ≤ 0.05. Groups not connected by a common letter are significantly different. * significant genotype effect, ** significant diet effect, *** significant genotype-diet interaction effect. C, control diet; FD, folate-deficient diet.

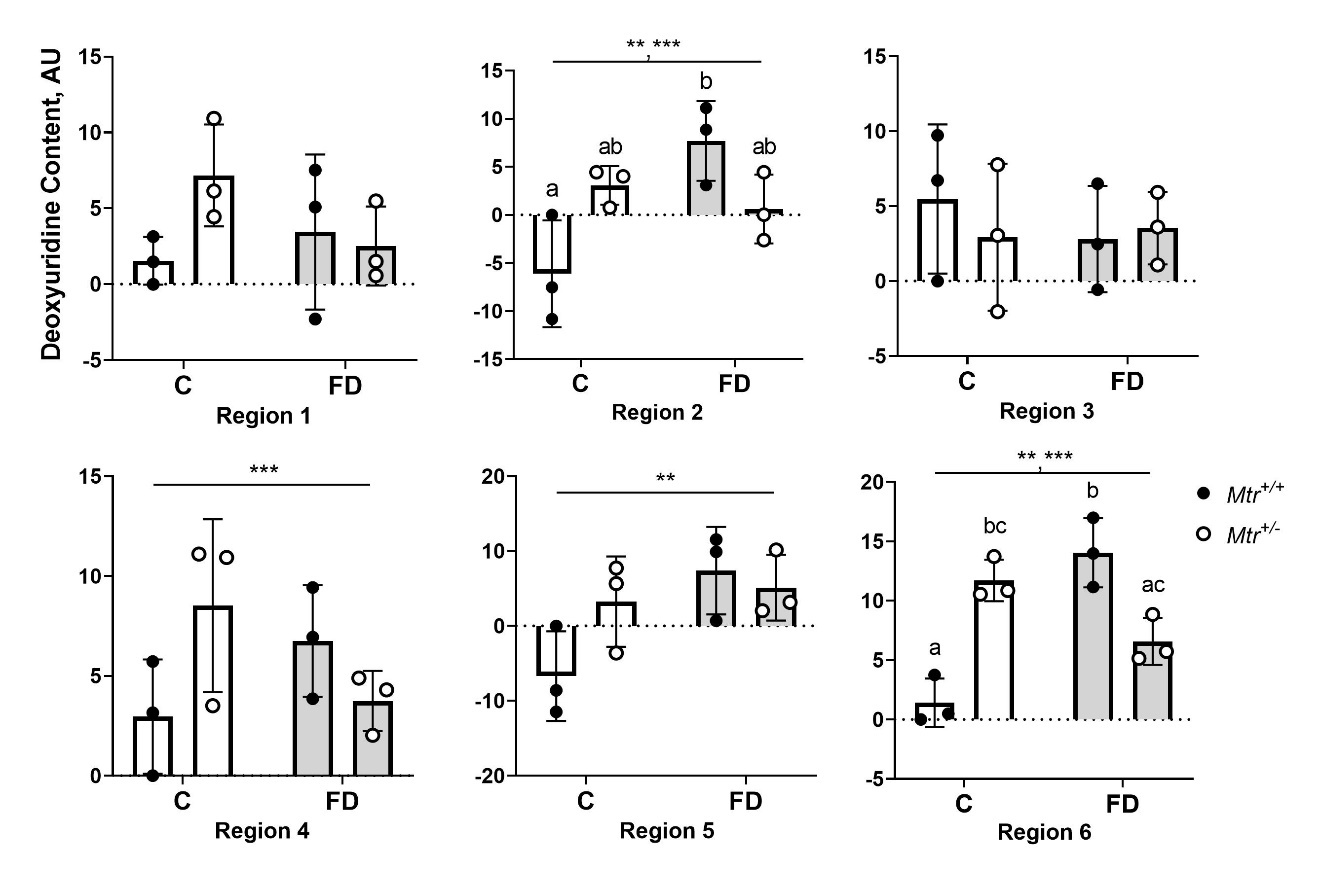

Figure S4. Uracil levels in liver mitochondrial DNA of *Mtr^+/+^* and *Mtr^+/-^* mice vary by mitochondrial genome region. Uracil was quantified by real-time PCR assay relative to wild type control sample; n = 3 per group. Data are shown as mean ± s.d. and were analyzed by two-way ANOVA with Tukey’s post-hoc analysis, with significance defined as *p* ≤ 0.05. Groups not connected by a common letter are significantly different. * significant genotype effect, ** significant diet effect, *** significant genotype-diet interaction effect. C, control diet; FD, folate-deficient diet; AU, arbitrary units.

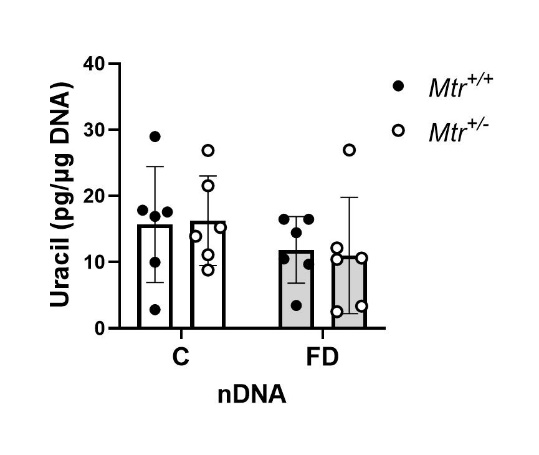

Figure S5. Uracil levels in nuclear DNA are not affected by *Mtr* genotype or folate deficiency in mouse liver. Uracil was quantified by GC-MS; n=6 per group. Data are shown as mean ± s.d. and were analyzed by two-way ANOVA with Tukey’s post-hoc analysis, with significance defined as *p* ≤ 0.05. Groups not connected by a common letter are significantly different. * significant genotype effect, ** significant diet effect, *** significant genotype-diet interaction effect. C, control diet; FD, folate-deficient diet; nDNA, nuclear DNA.
